# Distinct regulatory modules identified in the promoters of wheat *Glu-1* genes suggest different regulatory mechanisms

**DOI:** 10.1101/011635

**Authors:** Szabolcs Makai, László Tamás, Angéla Juhász

## Abstract

High molecular weight glutenin subunits of wheat are economically important seed storage proteins. They are coded by paralog pairs of the *Glu-1* gene on each of the three genomes in the hexaploid wheat. Their expressions are under both temporal and spatial control. Many factors have been identified that influence the activity of *Glu-1* genes, but the underlying regulatory mechanisms are still unclear. In order to identify motifs and motif clusters responsible for quantitative regulation of *Glu-1* gene expressions, promoter profiles and transcription dynamics of the genes were analysed. It was found that promoter motif compositions of homoeolog *Glu-1* genes are conserved. Our results demonstrated that while promoter profiles explain the differences of expression between homoeologs and between paralogs, it does not explain the variation of activity between alleles. Interestingly, our analyses revealed that the promoters of *Glu-1* genes are divided into six *cis*-regulatory modules that are either locally overrepresented by binding sites belonging to unique but distinct transcription factor (TF) families or have conserved motif clusters. Moreover, our analyses demonstrated that the varying expression dynamics of TFs across genotypes is likely to be the primary contributor of the allelic variation of *Glu-1* gene expressions. Thus, the six putative *cis*-regulatory modules in the *Glu-1* gene promoters bound by the differentially expressed TFs are suggested to play a key role in the quantitative and tissue specific regulation of these genes.

## INTRODUCTION

Wheat seed storage proteins (SSPs) are one of the primary sources of proteins in human diets and animal feed worldwide. These proteins are synthesized in the endosperm; a tissue specialized to starch and protein biosynthesis and storage. The prolamin superfamily of SSPs is a main component of wheat flour and their composition and ratio control dough properties, thus the quality of the end product. There are three main types of prolamin proteins: the sulphur (S) rich prolamins (alpha-, gamma-gliadins and low molecular weight (LMW) glutenins), sulphur poor prolamins (omega gliadins) and the high molecular weight (HMW) glutenins. Consequently, the quality of wheat dough is determined by the allele composition present in the genotypes and the proportion of the prolamin proteins thereby expressed.

The HMW glutenin subunits (HMW GS) of the hexaploid *Triticum aestivum* are encoded by 3 homoeologous loci denoted as Glu-A1, Glu-B1, Glu-D1, located on the long arm of chromosome 1 of all three genomes. As a result of a tandem duplication, the three Glu-1 loci encode two paralogs of the HMW glutenin subunit, called x and y-type or Glu-1-1 or Glu-1-2, making a total of 6 *Glu-1* genes present in the hexaploid wheat (Figure 1). Besides the two paralogs HMW glutenin genes, the Glu-1 locus also encodes for two paralog globulins, a receptor kinase and a serine/threonine protein kinase. The *Glu-1* genes are intronless, surrounded by transposable elements. The order and orientation of the genes is highly conserved in the loci across all three genomes (1). Earlier studies reported that ortholog genes show higher conservation than paralogs (2).

**Figure 1.**
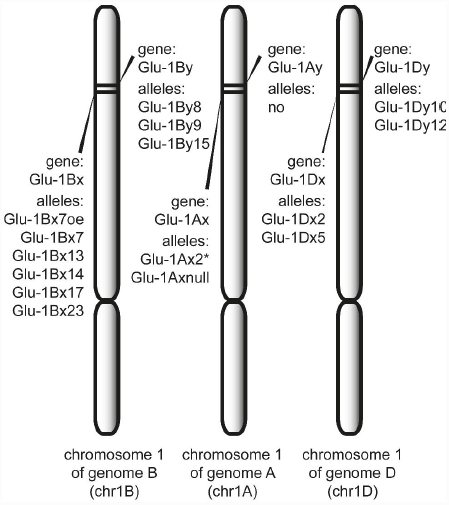
*Glu-1* genes are located at the long arm chromosome 1 of each of the three homoeolog genomes (B, A, D) of the hexaploid wheat. Alleles of each genes involved in this study are listed under their respective gene names.

HMW glutenin subunits are solely expressed in the endosperm; the 4-5 active genes in the hexaploid genome account for approximately 12% of the total seed protein content (3, 4). The expression levels of the homoeolog and paralog *Glu-1* genes vary greatly. In general, *Glu-1Bx* genes have the highest transcription level followed by *Glu-1Dx* genes. The y-type genes are the lowest expressed. *Glu-1Ax* gene has a null allele in most genotypes (5). It has been reported that the *Glu-1Ay* gene is always inactive in hexaploid wheat varieties (6). However active *Glu-1Ay* genes were identified from related species (7). Expression of the *Glu-1Ay* genes has been reported for hexaploid genotypes by Margiotta et al., 1996. This phenomenon is hypothesized to be the result of the introgression of the *Glu-1Ay* gene from *Triticum urartu* or *Triticum dicoccoides* species during the breeding programs (8). The inactivation of the *Glu-1Ay* is due to the gene disruption that varies between cultivars. In the cultivar Chinese Spring, a *Wis-3* retroelement insertion was found in the repetitive region of the HMW GS *Glu-1Ay* gene (9) while in cv. Cheyenne, an in-frame stop codon has been identified (10).

The synthesis of prolamin proteins is regulated mainly at the transcriptional level and is influenced by sulphur and nitrogen availability (11, 12). During grain filling, the accumulation of storage proteins is source-limited (13) which is in contrast with the sink-limited starch accumulation (14). Due to the linkage between the nitrogen and carbon metabolism, the endosperm is capable of absorbing excess nitrogen, as long as the N/C ratio is in equilibrium (15, 13). It has been proposed earlier that grain filling, as well as HMW GS synthesis, may be regulated at the plant level (16).

The expressions of all prolamin proteins follow a well characterized, although varying, temporal pattern during seed development. Their transcription is regulated by trans-acting factors associated with *cis*-acting elements as well as epigenetic factors (17–20). Earlier studies suggested that transcription of gliadin and LMW glutenin genes (*Gli-1,2* and *Glu-3*, respectively) were influenced by methylation and imprinting while expression of HMW GS coding *Glu-1* genes is less dependent on these epigenetic factors (19). A closer analysis of the expression profiles of prolamin genes and their responses to abiotic stresses indicate the presence of different regulatory mechanisms for each prolamin protein family (21, 22). A model for different transcriptional regulation mechanisms have already been proposed which is based on the conserved non-coding elements of the LMW GS coding *Glu-3* genes (20).

Earlier studies showed that promoters of the *Glu-1* genes have distinct features compared to the promoters of other proteins in the prolamin superfamily (23, 24, 17, 25, 26). Promoters of *Glu-3* genes as well as the sulphur rich prolamin genes of wheat or other monocots all have a bipartite conserved binding site (BS), called endosperm element. They consist of a DOF (DNA-Binding with One Finger) binding domain called Prolamin box (P-box) and a GCN4-like motif, also called the Nitrogen-box (N-box), that binds to a bZIP (basic leucine zipper) factor (27, 28, 25). This nitrogen related element has a role in down-regulation when nitrogen supply is low (29). However, *Glu-1* genes lack the N-box at -300. Instead, their conserved promoter regions include some specific motifs called CEREAL-box and HMW enhancer that contains similar sites to P-box (11, 30). The divergent expression of *Glu-3* gene types have been found to correlate well with their conserved non-coding regions (20). However, no similar study has been published yet for the *Glu-1* genes of HMW glutenin subunits.

There are many studies identifying transcription factors (TF) involved in the prolamin gene expression (31, 27, 32). Most of these TFs belong to the DOF, bZIP and MYB families. Additionally, NAC, LEC1 type TFs were also found to influence seed storage protein expressions (33, 34). In case of wheat, none of these trans-acting factors are exclusively expressed in the endosperm (35). The wheat prolamin binding factor (WPBF), a DOF type TF was reported to be constitutively expressed in all tissues. Indeed, many reports confirm that an interaction of these factors is the key to tissue specificity (32, 36, 37). Interaction of bZIP and DOF has been reported and this type of interaction appears to be unique to the endosperm of monocots (33). It was confirmed by Diaz and co-workers that barley GAMYB and DOF transcription factors were able to interact forming a complex, which activated endosperm specific gene expression (38).

HMW GSs have an important role in crop quality improvement programs. Considerably research has already been undertaken to identify and describe different alleles of *Glu-1*, because of their contribution to the formation of the gluten macro-polymer. However, it is equally important to assess alleles from a quantitative perspective. Knowing what influences the expression of *Glu-1* genes, understanding the underlying mechanisms of transcription, or identifying interacting TF proteins could further assist breeders in achieving better traits (39). In order to better understand the regulation of the genes of HMW glutenin subunits, the aim of our study is to characterize the promoter regions of *Glu-1* genes of wheat cultivars via an *in silico* analyses. A working model of combinatorial *cis*-regulation of *Glu-1* genes is proposed, which was formulated using a reverse approach as described in Werner and co-workers (40).

## METHODS AND MATERIALS

All publicly available gene sequences of HMW glutenin subunits of *Triticum aestivum* were collected from NCBI’s nucleotide archive. Additional promoter sequences for *Glu-1Ay2* and *Glu-Dx12* were downloaded from the wheat survey sequence repository (http://www.wheatgenome.org). Sequences were grouped by locus, paralog types (x or y), and genotypes. Altogether there were 156 HMW GS promoter sequences collected. 140 sequences were longer than 250 bp, 122 longer than 500 bp and 27 were longer than 700 bp. 87 promoters belonged to x-type and 69 to y-type HMW GS genes. The promoters represented well all three loci: there were 60 from the A genome, 67 from the B genome and 29 from the D genome. For sequence characterisation, promoters longer than 700 bp were used. List of motifs generally related to prolamin gene promoters were used based on the results of Juhász and co-workers. The terminology of motifs and “boxes” is the same as in the study of Juhász and co-workers (20). Additional motifs were used from PlantCare database. Promoter profiles were graphically represented using the Promoter Profiler tool developed by one of the authors. Locally overrepresented regions of binding sites on the promoter sequences were identified by statistical analysis. Promoters were screened for regions where either similar BSs are located at a distance of maximum 100 bp or where a BS has no other BSs located in its 50 bp long upstream or downstream region. Conserved motif clusters were searched in promoter sequences. Motifs clusters of less than 100 bp length that occurred at least half of the studied promoters were searched for following the framework CRÈME as described by Sharan and co-workers (41).

The expression analysis of HMW GS alleles were conducted *in silico* based on EST data and measured in transcript per million (TPM), as it was described earlier (20). TPM values were standardized to zero mean and unit deviation. Libraries used in the study are shown in Table 1. Parameters of similarity search and subsequent filtering varied by EST libraries and are shown in Table 1. Differential expressions of alleles across libraries were calculated by normalizing TPM data to a shared allele of the compared libraries.

**Table 1.** cDNA libraries of developing wheat seeds used in this study and their respective genotype information. Library referenced as DuPont has an unknown genotype and its allelic composition was inferred from best BLAST results. Setup for BLAST query and subsequent filtering are shown in parameters column (ws: word size, ug: ungapped penalty, pc: percent identity).

| Library ref. | Genotype | Allelic composition of HMW glutenin |  |  |  |  |  | Available libraries for developing seed (in DPA) | Number of ESTs | Source | BLAST Parameters |
| --- | --- | --- | --- | --- | --- | --- | --- | --- | --- | --- | --- |
|  |  | Ax | Ay | Bx | By | Dx | Dy |  |  |  |  |
| Chinese Spring | Chinese Spring | null | null | 7 | 8 | 2 | 12 | 5,10,20,30 | 10729, 11282, 11308, 12573 | NCBI dbEST IDs: 19555, 10469, 19547, 10479 | ws: 100, ug:30, pc:96 |
| Glenlea | Glenlea | 2* | null | 7oe | 8 | 5 | 10 | 5,15,25 | 5805, 5368, 5094 | TaGI: #A9H, #A9I, #A9J | ws: 100, ug:30, pc:96 |
| DuPont | unknown | 2* | null | 7 | 9 | 5 | 10 | 3,7,14,21,30 | 4689, 4887, 4967, 958, 840 | TaGI: #BSD, #BSE, #BSF, #BSG, #BSH | megablast, pc:96 |

Allelic composition of genotypes was collected using annotations supplied by their respective depositors or published results. The query sequences of each alleles used for EST based expression measurements are as follows: *Glu-1Ax2\** -M22208 (for Glenlea, DuPont libraries); *Glu-1Bx7* -DQ119142 (for Glenlea lib.) and BK006773 (for Chinese Spring and DuPont libs.); *Glu-1By8* -JF736014 (for the Chinese Spring and the Glenlea libraries); *Glu-1By9* -X61026 (for the DuPont library); *Glu-1Dx2* -BK006460 (for the Chinese Spring lib.); *Glu-1Dx5* – BK006458 (for the Glenlea and the DuPont libraries); *Glu-1Dy10* – X12929 (for the Glenlea and DuPont libraries); *Glu-1Dy12* – BK006459 (for the Chinese Spring library).

Expression of transcription factors families were calculated slightly differently because the query sequences were clusters and not experimentally isolated gene sequences. Therefore, expression was measured by TF types. Transcription factor cluster sequences were downloaded from Database of Wheat Transcription Factor (wDBTF) (42) and the Plant Transcription Factor Database (43). 102 bZIP sequences were used to measure the expression bZIP TF family, 28 for DOF, 103 for MYB-R2R3, 134 for NAC, 22 for NF-YB, 18 for NF-YC from these databases. In addition 87 AP2 and 15 VP1 TF sequences were downloaded from NCBI nucleotide database. Parameters of the expression measurements were as follows: megablast query and results filtering to percent identity 90, e-value cut-off 0.01.

Gene expression analysis under various environmental conditions was based on the published data of Hurkman and co-workers, where protein expression levels of seed grown under different temperature and nitrogen regimens were measured as spot volumes (22). This data was further analysed by our lab, where expression patterns of HMW GSs were studied individually. Spot volume data was first averaged by the three samples of each sampling time then their sum was calculated. Data were standardized to zero mean and unit deviation. Standardized values were hierarchically clustered by Cluster3 (44) using settings of centred absolute correlation and centroid linkage.

## RESULTS

### Motif composition of the 5’-UTR flanking region of *Glu-1* genes

Transcription factors bound to motifs of the 5’-flanking UTR of *Glu-1* genes are the main drivers of their transcription. Identifying binding sites and characterizing differences among the six *Glu-1* genes and their alleles offers a view on the underlying regulatory mechanisms. Since the allele composition of the six *Glu-1* genes are genotype specific, differences across alleles may be directly correlated with the phenotypes, thus with dough making quality, of the cultivars involved in this study.

The analysis of the motif composition of *Glu-1* genes showed that the proximal and distal promoter regions of the homoeolog genes are in general more similar than that of the paralogs (Figure 2 and 3). Similarities were observed for the pairs *Glu-1Ax/Dx* and *Glu-1Dy/By*. Promoter profiles of *Glu-1Bx* genes showed considerable differences compared to the other genes. Note that positions written in the text are within a range of ±10 bp for easier rading. (The complete list of motifs with exact positons for all studied sequences are in the supporting material.)

**Figure 2.**
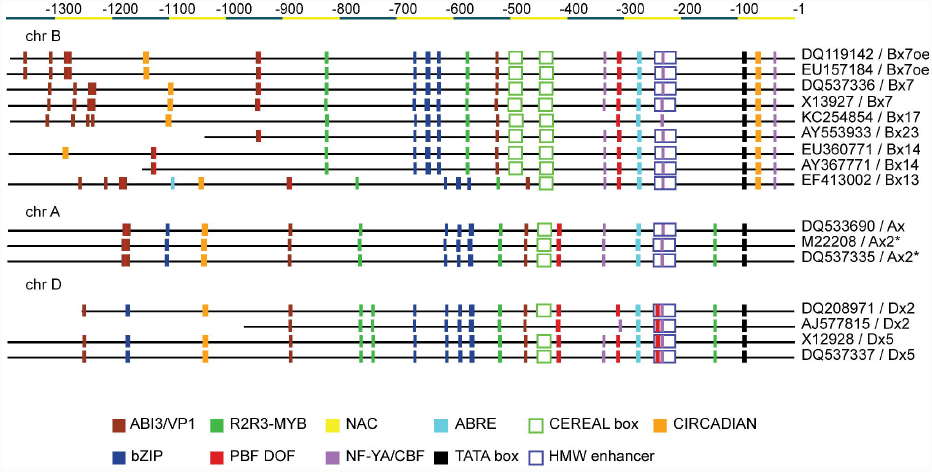
Promoter profiles of x-type HMW GS genes sorted by chromosomes. Only sequences longer than 700 bp are shown. Accessions and allele information are provided where available. Motifs in antisense direction are not shown but described in the text. Boxes are coloured by the TF family that is bound to binding site as described in Juhász et al., (2011). Promoter sequences are aligned to their TATA@92s because some promoters miss the 3’-end region. A colour legend is provided. Promoters show some shared as well as distinct features. The most outstanding promoter profile belongs to the *Glu-1Bx* genes that have duplicated CEREAL-boxes and a CIRCADIAN element at the proximal promoter region. *Glu-1Ax* and *Glu-1Dx* have more in common what is harmony with the phylogeny of hexaploid wheat.

**Figure 3.**
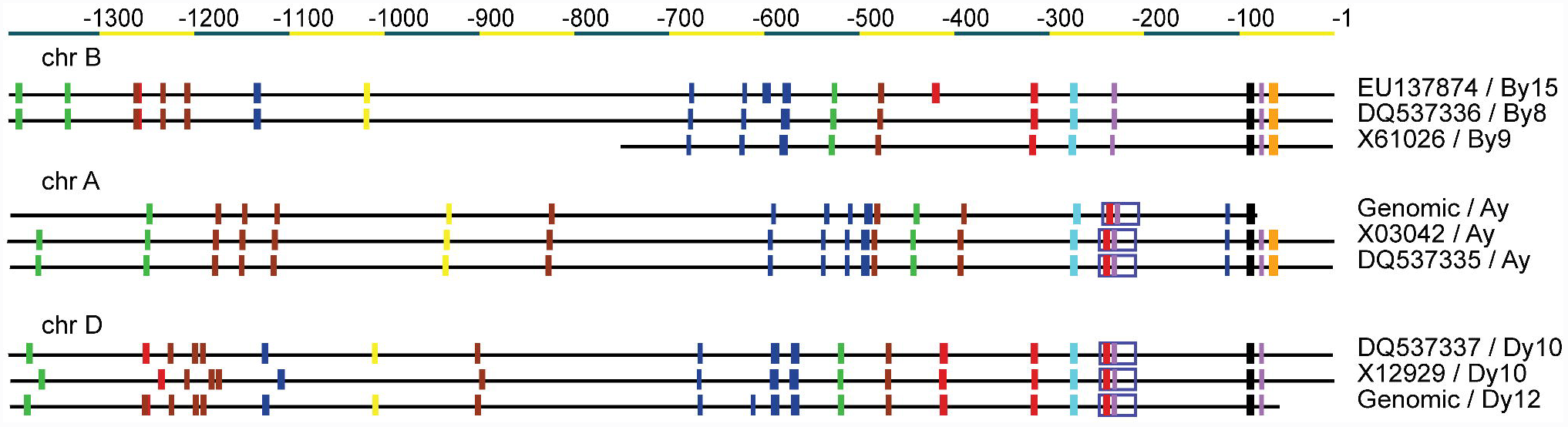
Promoter profiles of y-type HMW GS genes sorted by chromosomes. Only sequences longer than 700 bp are shown. Accessions and allele information are provided where available. Sequences retrieved from the wheat genome project are marked as “Genomic”. Motifs in antisense direction are not shown but described in the text. Colours of boxes are as in Figure 2. There distinct differences compared to the x-type *Glu-1* genes on Figure 2. The y-type *Glu-1* genes also have a promoter profile that reflects the phylogenetic history of hexaploid wheat. *Glu-1Dy* and *Glu-Ay* genes are more related than *Glu-1By.* No intact HMW-enhancer element could have been detected on *Glu-1By* genes, however its core palindrome site and a CCAAT box is remained.

The greatest difference between the analysed promoter sequences occurred in a region between the ABRE motif at -277 bp relative to transcription starting site (TSS) in sense direction (referred to as ABRE@277s where ‘s’ is for sense and ‘as’ is for antisense’) and the AACA/TA motif positioned between -400 and -500. Exact positions varied for the different alleles. In the case of *Glu-1Bx* genes (with the exception of *Glu-1Bx13*), there is an 55 bp long insertion that resulted in a duplicated CEREAL-box and the loss of a P-box at -418. In the case of *Glu-1Ay* genes, there is a 131 bp long deletion resulting in the loss of CEREAL-box and the loss of P-box at -300 bp. In the same region at around -312 bp, *Glu-1Ax* genes also lack this P-box.

*TATA boxes.* The TATA-box was always found at -92 relative TSS in all sequences analysed.

*CEREAL-box.* All x-type genes have a CEREAL-box at position -450 bp as shown in Figure 2 with polymorphisms characteristic to the genomes of hexaploid wheat. All *Glu-1Dx* genes have a single CEREAL box with a sequence of 5’-GACATGCTTAGAAGCTTTGAGTG-3’, whereas the CEREAL-box identified in *Glu-1Ax* type genes differs in a single nucleotide polymorphism (SNP): 5’-GACATGCTTAGAAGC<u>**A**</u>TTGAGTG-3’ (SNP is marked with **bold** and <u>underlined</u>). However, promoters of *Glu-1Bx* type genes usually have a double CEREAL box at -502 and -448, with a sequence of 5’-GACATG<u>**G**</u>TTAGAAG<u>**T**</u>TTTGAGTG-3’ that contains two SNPs which makes a 3’-CCAAT-5’ site on the antisense strand. The region between the two CEREAL-boxes is identical for all *Glu-1Bx7* genes.

*HMW enhancer.* A general feature of *Glu-1* gene promoters is the so called HMW enhancer element which is present in almost all analysed promoter regions except the *Glu-1By* type alleles. Three variants of HMW enhancer motifs were detected at a well conserved location of -247. *Glu-1Ay* and both x-type and y-type Glu*-*1D genes have a sequence of 5’-GTTTTGCAAAGCTCCAATTGCTCCTTGCTTATCCAGCT-3’. *Glu-1Ax* genes have an SNP in the sixth position (GTTTT<u>**A**</u>CAAAGCTCCAATTGCTCCTTGCTTATCCAGCT). *Glu-1Bx* genes have a variant with one deletion and two SNPs as follows: 5’-GTTTTGCAA-GC<u>**A**</u>CCAATTGCTCCTT<u>**A**</u>CTTATCCAGCT –3’. Glu-1Ax, Glu-1Dx and Glu-1Dy genes have an additional partial HMW enhancer element at -417, -420 and -418, respectively, keeping only its core palindrome site: 5’-TTTGCAAA-3’. Promoters of the analysed *Glu-1By* genes also possess this partial HMW enhancer element.

*Prolamin Binding Factor (PBF DOF) binding sites.* Sequences were searched for motifs of P-box to identify possible PBF DOF binding sites. In the case of *Glu-1Dx*, -*1Dy* and -*1Ay* genes there is a P-box (TGCAAAG) at -243 as part of the HMW enhancer. *Glu-1Ax*, *Glu1-Dx* and *Glu-1Dy* genes and the *By15* allele of *Glu-1By* genes have a P-box in a conserved position at -418 bp. *Glu-1Dx*, *Glu-1Dy, Glu-1Bx* and *Glu-1By* genes have an occurrence of P-box at -312 bp. *Glu-1Bx* genes have an additional P-box at around -945. *Glu-1Bx* and *Glu-1Ax* have a distal P-box occurrence between 1100-1200 bp, exact positions vary on insertions and deletions described above.

*bZIP binding sites.* With the exception of *Glu-1Ay* promoters, all genes have a double N-box (TGAGTCA) with a distance of 21 bp. Their positions vary between -550 and -650 bp. *Glu-1Ay* genes have a single N-box at -413. No N-box was detected on the antisense strand.

Skn-1 like motif (GTCAT) has been searched on sequences. *Glu-1Ax* and *–1Bx* genes have two occurrences of Skn-1 on the sense strand at the positions -569, -615 and -645, -670, respectively. *Glu-1Dx* and *-1Dy* have three occurrences in sense direction. Two of these positions are conserved at around -568 and -670. The third is placed between these two at -613 and -590 depending on alleles. With the exception of *Glu-1Ay* genes all, *Glu-1* promoters have a Skn-1 like motif at -586 in antisense direction. *Glu-1Bx*, *Glu-1By* and *Glu-1Dx* genes have an additional motif at around -730 in antisense directions, although exact position depends on insertions and deletions.

*Glu-1Ay* genes have the most occurrences of Skn-1 like element. Five occurrences are in sense direction and one at antisense. Their positions are -512, -456, -431, -410 and -115 on sense and -625 in antisense direction.

*MYB binding sites.* All analysed promoters have an AACA/TA motif at position -520 (-578 for genes encoded in the Glu-B1 locus), which in the case of *Glu-1Ax* genes is preceded by a CAAT element. Only the x-type *Glu-1* gene promoters have a further AACA/TA motif at -172 in antisense and at -737 in sense direction. According to the analysed sequences, *Glu-1Dx* and *-1Ax* promoter regions have an additional AACA/TA motif at position -142. There is a unique MYB1AT (5’-AAACCA-3’) site at – 743 in all promoters of *Glu-1Dx* genes. Unique to *Glu-1Ay* and *Glu-1Dy* is a MYB1AT motif at -685 in antisense direction. *Glu-1Bx* genes have an additional MYB1AT at -548 in antisense direction.

*VP1 binding sites.* The promoters of the *Glu-1* genes have an ABRE motif (ACGTGGC) at -277. In addition, all analysed *Glu-1* gene promoters have a RY core site element (CATGCA) at around -470. *Glu-1Ay* genes have an additional RY core site downstream at position -396. In the 2000 bp long 5’ flanking region it occurred only once for all genes except *Glu-1Ay*. *Glu-1Ay* has an additional ABRE motif at -864 bp. Promoters above -700 bp are rich in RY core site elements (data not shown).

*NF-YA (CBF or LEC1) binding sites*. Transcription factors belonging to this family is bound the CCAAT box, therefore they are also called the CCAAT-box binding factor (CBF). The CBF@237s is well conserved in all analysed HMW GS genes. *Glu-1* genes of the B genome have additional CCAAT boxes at -36 and -336. A further CCAAT box was identified in *Glu-1Bx* genes at -642 in antisense direction. *Glu-1Ay* and *Glu-1Dy* have one extra motif in antisense direction at the -511 and -585 positions. There is one further CCAAT box between -600 and -1000 for all analysed genes with the exception of *Glu-1Ax* which has two occurrences in the region.

*Additional motifs.* All *Glu-1Bx and -1By* genes have a CIRCADIAN motif (CAATCTCATC) at around -69 and all x-type *Glu-1* have an additional motif around at -1140. Above -700 bp, the motifs are apparently less frequent and less conserved. However, while the motif occurrences of this distal upstream region are relatively conserved, their positions are more polymorphic due to insertions and deletions described above (data not shown). Promoters of *Glu-1* genes have multiple NAC binding sites above -900, while no occurrence was detected for x-type genes.

The identification of BSs on the promoter region of *Glu-1* genes showed a high conservation across similar genes and variation across the six *Glu-1* genes.

### Identifying cis-regulatory modules

Determining the positions of single BSs is necessary but not sufficient to “decode” the regulatory mechanisms programmed in the promoters of the *Glu-1* genes. Therefore an analysis to determine the distribution, local overrepresentation and clusters of binding sites was carried out. In conclusion, during this analysis, we found that single BSs and certain motif clusters follow a highly conserved, non-overlapping distribution. Consequently, the promoters of *Glu-1* genes can be divided into six distinct *cis*-regulatory moduls (CRM) (Figure 4).

**Figure 4.**
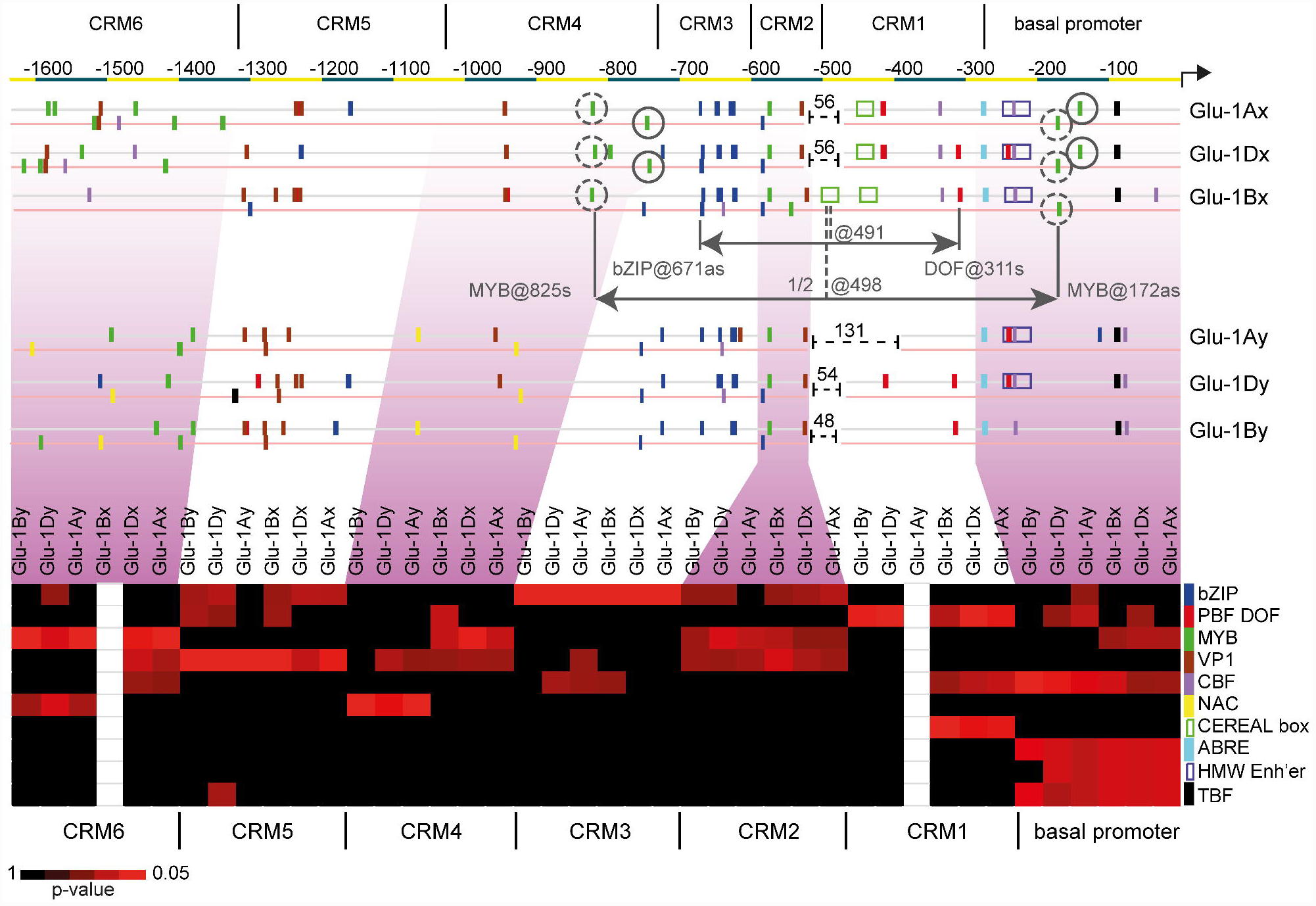
*Cis*-regulatory modules of the 1600 bps long 5’-UTR of *Glu-1* genes by locus. Colours of binding sites are as in Figure 2. Genes are grouped by x and y type. Sense and antisense strand is shown in grey and in light red, respectively. Gaps were inserted to the sequences between the CRM1 and CRM2 regions to align the highly conserved motif cluster of the CRM2. The insertions are marked with dashed lines and they lengths are shown. Complementary pairs of MYB BSs are encircled. Dashed and continuous circles represent separate pairs. The geometric distance and centre position of MYB BS pair are drawn for *Glu-1Bx* genes as example. Also for *Glu-1Bx* gene, a pair of DOF and bZIP BS is shown and its distance and centre position is marked. Interestingly, both centre positions are situated at the site of CEREAL@501s. Without the inserted gaps, *Glu-1Ax* and *Glu-1Dx* genes have similar structures. Heat map represents the p-values of the local overrepresentation by modules, BS types and genes. *Glu-1Ay* gene lacks CRM1 region and *Glu-1Bx* gene has no CRM6 region. The six CRMs and the basal promoter region show distinct features across genes.

In the 1600 bp long 5’-UTR region of *Glu-1* genes, there were four regulatory regions identified that are locally overrepresented by binding sites belonging to a single transcription factor family. Three additional regulatory regions were found to have conserved motif clusters. These modules have a conserved order and oriantation for all genes. CRM1 is overrepresented by DOF, CRM3 by bZIP, CRM5 by VP1 and CRM6 by MYB binding sites. The CRM1 is situated between -490 and -360 bp and has 1 to 2 DOF BSs. CRM3 was identified between -750 and -610 bp and it harbours 3 to 5 bZIP recognition sites. CRM5 is localized between -1330 and -1230, and CRM6 between -1650 and -1340. The number of BSs in CRMs are characteristic to genes. *Glu-1Ax*, *Glu-1Bx* and *Glu-1By* have one DOF BS, while the *Glu-1D* genes have two in CRM1 region. *Glu-1Bx* and *Glu-1Ax* have two bZIP BSs while *Glu-1D* and *Glu-1By* genes have three in their CRM3 regions. *Glu-1Ay* have five bZIP BSs in CRM3. CRM5 and CRM6 regions have a motif compositions conserved across orthologs. Interestingly, *Glu-1Ay* gene has no CRM1 region, and *Glu-1Bx* gene completely misses the MYB BS abundant CRM6 region.

Motif clusters, where at least two BSs are in combination in conserved position and orientation were identified. CRM2, CRM4 and the 277 bp long proximal promoter of *Glu-1* genes region have such conserved motif clusters. The proximal promoter region of *Glu-1* genes streches upto the ABRE@277s and we will refer to this as the basal promoter region to express its basic role in the transcription of *Glu-1* genes. It contains a conserved composition of TATA@92s, CBF@37s, the HMW-enhancer and the ABRE@277s. It has additional MYB BSs for x-type *Glu-1* genes (see description later). CRM2 is the only module that has a motif cluster conserved across all *Glu-1* genes. It has one VP1 BS and a MYB BS at a distance of 45-50 bp. In the case of *Glu-1Bx* genes, there is an additional MYB@548as in the CRM2. The CRM4 of x-type *Glu-1* genes have several MYB BSs in conserved positions, while CRM4 of y-type *Glu-1* genes have NAC BSs also in a conserved manner.

Remarkably, the number of MYB BS in CRM4 and the in basal promoter region are correlating. For *Glu-1Ax* genes, the basal promoter region has two MYB BSs and the CRM4 region has also two MYB BSs at conserved locations. Similarly, *Glu-1Dx* genes have two MYB BSs in the basal promoter region and have 3 at its CRM4 region, all at conserved locations. *Glu-1Bx* genes have one at each regions, also at conserved locations across all alleles/genotypes involved in this study. These MYB BSs appear to be conserved in remote, complementary pairs, in other words, a MYB BS on the sense direction has a MYB BS pair on the antisense strand. These pairs are as follows: MYB@825s-MYB@172as for *Glu-1Bx* genes, MYB@768s-MYB@174as and MYB@692as-MYB@143s for Glu-1Ax genes and MYB@764s-MYB@173as and MYB@737as-MYB@142s for *Glu-1Dx* (Figure 4). Interestingly, in the case of y-type HMW GS, there are no MYB BSs detected in either regions.

The 5’-end of the coding region was also searched for motifs. In the case of *Glu-1Bx* genes, we found a conserved motif cluster of MYB – bZIP BSs at position (+) 238 bp at a distance of 73 bp (data not shown). All other genes involved in our study had no conserved motif cluster in the 5’-end of the coding region.

We identified six *cis*-regulatory modules and a basal promoter region in the 1600 bps long 5’ flanking sequences of *Glu-1* genes as shown in Figure 4. While these CRMs vary across genes, they seem to be conserved across alleles, thus across genotypes.

### Expression profiles of transcription factors binding to different CRM regions

*Glu-1* genes are expressed solely in the endosperm of wheat seeds. Transcription factors interacting with its CRM and the basal promoter regions play a major role in their quantitative regulation. In order to understand how and when these interactions happen, an expression analysis of interacting TFs was carried out in developing seed. Three genotypes were selected for our EST based transcription analysis where there were reliable EST data available for developing seeds of hexaploid wheat (Figure 5A and Table 1). Transcription factors binding to the CRMs of *Glu-1* genes belong to the following TF families: DOF, bZIP, VP1/RY, AP2, NAC and MYB. Due to the lack of a comprehensive list of wheat TFs, we used publically available TF gene clusters and our analysis was restricted to measure expression dynamics of the above mentioned TF families as opposed to expression dynamics of single genes. However, since our goal was to describe tendencies rather than identify genes, this did not present a constraint to the analysis.

**Figure 5.**
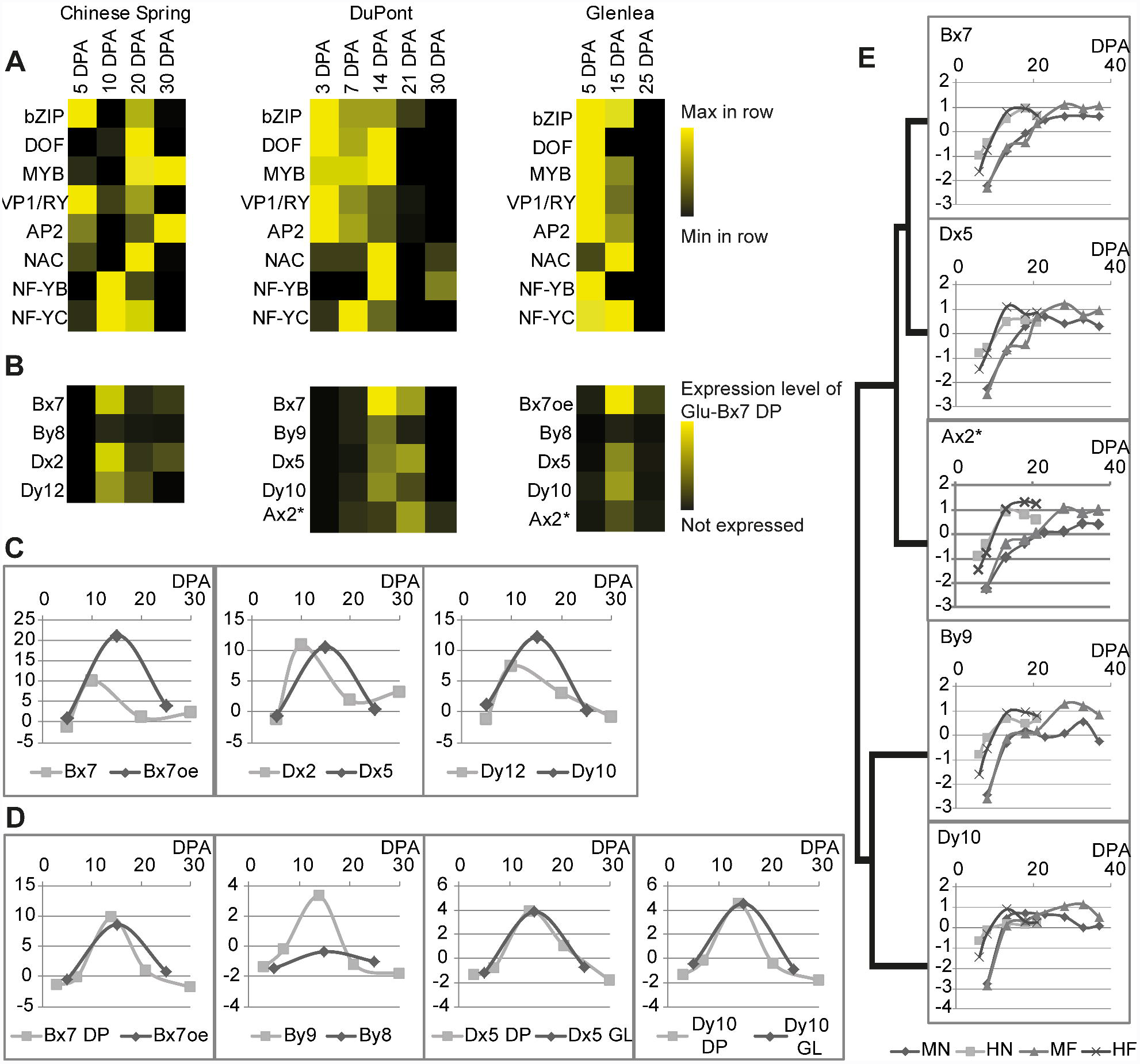
Expression profiles of transcription factor families interacting with the CRMs of *Glu-1* promoters (A) and the expression profiles of *Glu-*1 genes in the same libraries (B). Expression values of TF families were standardized to zero mean and unit deviation by rows. Expression data of *Glu-1* genes are standardized to zero mean and to unit deviation by libraries. TF expression dynamics of the DuPont and Glenlea libraries have similar patterns. Expression dynamics of *Glu-1* genes of the DuPont and Glenlea libraries also demonstrated similar patterns. *A*lleles of *Glu-1* genes were compared across the different genotypes (C-D). Data normalized to *Glu-By8* of the Chinese Spring and the Glenlea libraries are shown to demonstrate differences between alleles of *Glu-1Bx*, *Glu-1Dx* and *Glu-1Dy* genes (C). While *Glu-1Dx2* and *Glu-1Dx5* have similar level of activity, *Glu-1Dy10* had higher activity than *Glu-1Dy12*. As expected, the overexpressing *Glu-1Bx7oe* of Glenlea had higher activity than the normal expressing *Glu-Bx7* of Chinese Spring. Expression data of the DuPont libraries and Glenlea were also cross-examined by normalizing expression data to *Glu-1Ax2\** (D). Alleles of *Glu-1Dx* and *Glu-1Dy* genes showed similar profiles as well as *Glu-1Bx7oe* and *Glu-1Bx7 DP*. The only difference was detected between the *Glu-1By* genes. *Glu-1By9* had higher activity than *Glu-1By8*. Expression profiles of proteins under various growing conditions are shown in E. A tree was constructed by hierarchical clustering based on the standardized expression values. Key to treatments: MN: moderate temperature without fertilizer, MF: moderate temperature with fertilizer, HN: high temperature without fertilizer, HF: high temperature with fertilizer. Genes of x and y types give different responses to abiotic changes. The response to N (MF) gave the most distinct response between the genes. The axis X represents DPA in all diagrams (C-E).

The three genotypes had varying expression dynamics for the analysed TF families. The greatest difference in expression dynamics across genotypes was detected for DOF type TFs. In the Glenlea library, DOF TFs have a peak at 5 DPA while in the Chinese Spring library, DOF TFs have a peak at 20 DPA and no expression levels were detected at 5 DPA. In the DuPont library, the DOF expression level is peaked at 14 DPA but was present as early as 3 DPA. No expression was detected at 21 and 30 DPA.

NAC type TFs also showed diverse expression dynamics. While the Glenlea libraries gave an expression peaking at 15 DPA, both the Chinese Spring and the DuPont libraries had a profile with peaks at 20 and 30 DPA, respectively. Nevertheless, in case of the DuPont library the second peak could be an artefact as the number of EST data for 21 and 30 DP are one fifth of the number of ESTs for 3, 7 and 14 DPA (Table 1).

The analysed MYB-R2R3 type TFs had similar expression curves for the Chinese Spring and the DuPont libraries with a peak at 5 DPA and a second, higher plateau at 20 DPA. However, MYB-R2R3-type transcription factors showed an opposite expression profile in the Glenlea library, as expression levels decreases with time according to EST data.

Transcription factors belonging to bZIP family showed similar expression dynamics. All three genotypes have an early peak with a varying but maintained level until 20, 14 and 15 DPA for the Chinese Spring, the DuPont and the Glenlea libraries, respectively. AP2 and VP1/RY had an early peak at around 5 DPA in all studied genotypes.

The analysis demonstrated that the transcription activity of TFs binding to CRMs of wheat *Glu-1* genes varied across the three genotypes. However the expression profile of the DuPont library and the Glenlea library had higher similarity compared to the Chinese Spring library.

### Expressions of Glu-1 alleles

In order to see how the variation of transcription factor expressions profiles influence the expression of their target, *Glu-1* genes, an expression analysis of *Glu-1* genes was also carried out on the same EST libraries. Alleles of the *Glu-1* genes were collected from online databases for all three genotypes as seen in Table 1.

The analysis demonstrated varying expression profiles across the analysed genotypes (Figure 5B). In the Chinese Spring cDNA library, transcripts of *Glu-1Bx7* and *Glu*-*1Dx2* alleles have an early peak at 10 days post anthesis followed by a decrease at 20 DPA. At 30 DPA, it turned around as there was an increase at 30 DPA in the expressions of *Glu-1Bx* and *Glu-1Dx* genes. Unfortunately, no data was available beyond 30 DPA. Expressions of *Glu-1Dy* and *Glu-1By* genes were less correlated. *Glu-1Dy12* gene has a distinct transcription profile and after a peak at 10 DPA a constant decline in activity was demonstrated at 20 and 30 DPA without the turning point during the studied period. Transcription of *Glu-1By8* gene was maintained at low level all the way through of the studied period of seed development.

In the Glenlea library, there was a higher similarity in expression levels and dynamics between *Glu-1Dx* and *Glu-1Dy* genes (Figure 5B) compared to those in the Chinese Spring library. Additionally, the *Glu-1Bx* gene showed different expression dynamics. After standardizing TPM values, *Glu-1Bx7oe* of Glenlea demonstrated a 1.4x higher activity compared to *Glu-1Bx7* present in cv. Chinese Spring. Similarly, normalizing to the shared allele of *Glu-1By8* revealed that *Glu-1Dy10* was more active than *Glu-1Dy12* (Figure 5C). In the Glenlea library, y-type genes had always lower activity than x-type genes.

*Glu-1Bx7oe* allele is the highest expressing gene while the *Glu-1By8* allele is the least expressed one. The transcript level of *Glu-1Ax2\** allele is more than double that of *Glu-1By8*. The *Glenlea* libraries contain data up to 25 DPA. Consequently, the presence of a turning point similar to *Glu-1Bx7* allele in Chinese Spring libraries could not be confirmed. Nevertheless the expression of *Glu-1By8* allele is similarly low for both cultivars.

The early dynamics and transcript levels of HMW glutenin subunit genes from the DuPont libraries drew similar patterns when compared with the genes of cv. Glenlea. The only difference in expression was that for *Glu-1By* genes. The *Glu-1By9* allele from the DuPont libraries revealed a higher expression level than that of *Glu-1By8* after normalizing expression values to the shared allele of *Glu-1Ax2\** (Figure 5D). All other genes demonstrated similar activity, including the *Glu-1Bx* genes of DuPont and Glenlea.

In all libraries, x-type genes had, on average, higher activity than y-types. The expression dynamics of *Glu-1* genes in the Glenlea and the DuPont libraries were similar just like in the case of TF expressions. Differences between alleles were presented and this demonstrates that genes with identical promoter profiles have diverse expression activity. Consequently, the characteristic differences of expression dynamics of *Glu-1* paralogs together with their distinct motif compositions of CRMs suggest that there may be more than one regulatory mechanism involved in the control of *Glu-1* genes.

### Environmental effect on the expression of *Glu-1* genes

Differential responses to environmental changes can highlight different regulatory mechanisms. Therefore, to detect multiple regulatory mechanisms at action to control of *Glu-1* genes, an analysis of expression dynamics of *Glu-1* genes under abiotic stress was carried out. This analysis was using protein gel spot volume data of HMW GS published by Hurkman and co-workers (22). Wheat was grown under four different regimes after flowering. In short, the treatments were as follows: added fertilizer/nitrogen (MF), elevated temperature only (HN), both fertilizer and temperature (HF) and a normal, control environment (MN). Expression data were standardized and presented by genes in Figure 5E.

Elevated heat had an accelerating effect on the kinetics of protein synthesis which resulted in expressed HMW glutenin subunits present as early as 6 DPA. Added N gave different responses to the gene expression levels. A temporary fall back of expression (plateau in the curve) of *Glu-1Bx7* and *Glu-1Dx5* was detected between 10 and 20 DPA under added nitrogen treatment while *Glu*-*1By9* and *Glu-1Dy10* showed no response until 20 DPA than they increased their activity. Although the expressed volume of *Glu-1Ax2\** increased, it also had a temporary plateau between 10 and 20 DPA. *Glu-1Bx7* had no additional response to extra nitrogen compared to its response on elevated temperature, while expression levels of all other genes were further increased. All proteins were increased at late stages as a response to added nitrogen. Based on standardized data a hierarchical clustering was carried out to find correlation between expression patterns. The resulted tree (Figure 5E) have two main branches representing the paralog, x and y type genes. However, the expression of *Glu-1Bx7* and *Glu-1Dx5* are more related than that of *Glu-1Dy10* and *Glu-1By9* genes.

These data revealed that signals related to extra nitrogen at early to mid-developmental stage caused a temporarily decreasing effect on x-types genes and seemingly no effect on y-type genes. This temporary effect occurred at the same time when the transcriptional turning points of x-type Glu-1 genes were detected.

## DISCUSSION

In order to identify motifs and motif compositions responsible for quantitative regulation of HMW glutenin subunit expression in hexaploid wheat, promoter profiles and transcription dynamics of the *Glu-1* genes were analysed. It was found that promoter motif compositions of the same gene (e.g. *Glu-1Ax* genes or *Glu-1Dy* genes, etc.) are conserved and they do not reflect allelic variation. Our results confirmed that promoters of the homoeolog *Glu-1* genes are more similar to each other than the paralogs (2). While promoter profiles explains the differences of expression across the genes of different loci and of different *Glu-1* gene types (x and y), it does not explain expression variation between different alleles as it was assumed for LMW GS (*Glu-3*) genes in our earlier study (20).

The key finding of this study is the modular distribution of transcription factor binding sites on the promoters of *Glu-1* genes. Their 5’ UTR region can be divided into six *cis-*regulatory modules (CRM) and a basal promoter region. These regions are either locally overrepresented by binding sites (BSs) belonging to single TF families or have a motif cluster of highly conserved order and orientation (Figure 4). This structure is in sharp contrast with the diverse composition of the conserved non-coding regions of *Glu-3* genes where MYB, bZIP and DOF BSs are always in the proximity of each other. There are multiple DOF BSs around the positions -400 and -300 of *Glu-3* promoters similar to the region in *Glu-1* gene promoters where the CRM1 region is situated, while in the case of *Glu-3* these sites have two or three adjacent bZIPs (20). Similarly, at the location where *Glu-1* genes have a bZIP abundant CRM3 region (around -580), *Glu-3* genes also have multiple bZIP binding sites but surrounded by DOF BSs. The bZIP binding N-box on the *Glu-1* gene promoters was described as a suppressor in case of low N-supply (29). N regulation of *Glu-3* is controlled by the transcription factor complex that involves the DOF type wheat prolamin-box binding factor (wtPBF) (45). This difference indicates that N dependence is “wired” differently for the genes of LMW GSs compared to that of HMW GSs.

Further important difference between *Glu-1* and *Glu-3* genes concerning their regulation is that the higher molecular weight glutenin genes are not under epigenetic control. Consequently, tissue specificity of *Glu-1* is likely to be coded in their *cis*-regulatory elements (19). The modular distribution of motifs on *Glu-1* promoters can function similarly to *cis*-regulatory modules as described in the case of the *endo16* gene of sea urchin embryo (46). Istrail and Davidson demonstrated that different modules contribute differently to the expression of *endo16* and their combinatorial effect insures tissue specificity and the special expression kinetics during development (47). Inspired by this model, we assume that the identified CRMs control tissue specificity and expression dynamics of the *Glu-1* genes in a co-operating manner.

The single most conserved module is CRM2 that is present in all genes showing a conserved order and orientation of BSs. Indeed, it seems that all insertions and deletions of the promoter occurred upstream or downstream of this area, which indicates an essential role of CRM2 in the regulatory mechanism. Bioinformatic analysis have already demonstrated that tripartite elements of bZIP, MYB and RY-elements (VP1) are evolutionary conserved and appear to synergistically contribute to auxin-inducible expression (48).

The variance of the two most *Glu-1* specific motifs, namely the CEREAL-box and the HMW enhancer element was the most divert single elements between x and y type genes. Both elements are located in the CRM1 region. The promoters of y-type *Glu-1* show wider polymorphism at their HMW-enhancer sites than x-type *Glu-1* promoters. *Glu-1By* genes only have a partial HMW enhancer keeping a CBF element. The D genome has a variation of the HMW enhancer sequence including an additional P-box. As for CEREAL-boxes, their number seems to be directly proportional to expression activity. *Glu-1Bx* genes have two CEREAL-boxes, and they demonstrated the highest transcriptional activity. They are followed by all other x-type genes carrying only one CEREAL-box in the promoter region. The y-type *Glu-1* genes have the lowest activity and they lack CEREAL-box. In the case of *Glu-1Ay* gene, the complete CRM1 region is missing, which may be related to its inactivity.

CRM1, CRM3, CRM5 and CRM6 are the regions locally overrepresented by a BS belonging to a single TF family. However there are differences in the number of BSs within these regions. The number of motif occurrences on promoter sequences of *Glu-1* genes are related to expression levels as it was proposed by Chiu and colleagues (2012) in their published mathematical model (49). Nevertheless in wheat, the highly expressed *Glu-1Bx* genes have one single DOF binding P-box motif while *Glu-1Dx* and *Glu-1Dy* have two DOF binding motifs in CRM1, yet they are less expressed. Consequently, since most TFs present in the seed do not follow the expression dynamics of storage proteins, simple stoichiometric ratio of binding sites and spatial, temporal co-presence of transcription factors could result in differences in the expression levels. In the case of *Glu-1Dx* and *Glu-1Dy* genes, TF complex formation may take longer because there are several DOF binding sites competing for the same number of available transcription factors. For *Glu-1Bx* gene, transcription can be more effective for the same reason, as it has no self-competing BSs in its CRM1 regions and transcription factor complexes can be assembled faster.

CRM4, CRM6 and the basal promoter have motif clusters typical to either the x-or y-type *Glu-1* genes. The y-types genes have NAC BSs in their CRM4 and CRM6 while x-type genes do not have these BSs at all. It was already suggested that ENAC1, a NAC transcription factor, can be involved in seed development and abiotic stress response in rice (50). As for the x-type *Glu-1* genes, we found a highly conserved pattern of MYB BSs in the CRM4 and in the basal promoter region. The importance of the interaction of MYB TFs is well known for plants. In the case of *Arabidopsis thaliana*, it was demonstrated that an interplay of the MYB-R2R3 TFs controls the transcription of aliphatic glucosinolate genes (51). An interesting observation is that CEREAL-box is located at the geometric center position of the remote, complementary pairs of MBY BS of the CRM4 and the basal promoter regions (Figure 4).

Earlier studies showed that promoter region upto -277 nucleotides of *Glu-1Dx5* is enough to secure tissue specific expression of the transgene *uidA* but at a low level (52, 53, 23). This suggests a basic role of the 277 long basal promoter regions of *Glu-1* genes. It has already been proved that DOF, MYB and bZIP TFs of Tobacco are “compatible” with wheat promoters (54). Consequently, the fact that transcription factors are interchangeable demonstrates that the regulatory logic is conserved across organisms. This indicates that interaction and combination, or in other words the position and the synchronicity of binding events ensure tissue specificity rather than single tissue specific TFs (40).

Based on the *cis*-regulatory modules of the promoters of *Glu-1* genes identified in our study and known transcription factor interaction involving binding sites of CRMs, a combinatory *cis-*regulatory model is proposed (Figure 6). The tissue specificity of *Glu-1* genes is maintained by the basal promoter region via suppression. Factors bound to the HMW enhancer (or to the CCAAT-box in it) and to ABRE motif at -277 bp suppress transcription in all tissues but the endosperm. The CCAAT-box of HMW enhancer binds NF-YA TF which in combination with NF-YB and NF-YC TFs have an important role of combinatory regulation in plants (55). At the initial phase of endosperm development, the lack of a single component is sufficient to cease inhibition and a basic, low level transcription starts. The CRM1 and CRM3 modules are interacting and act as transcription enhancers. Their communication is likely to be mediated by the highly conserved CRM2 module that contains a tripartite element and requires the right combination of MYB, bZIP and VP1 TFs. It is hypothesized that a DNA loop is formed by a TF complex at CRM2 and that brings CRM3 and CRM1 in proximity. This is supported by earlier studies reporting that interaction between MYB and bZIP TFs can form DNA loops at a relatively short (>150 bp) distance (56). Once, PBF DOFs emerge in the endosperm tissue and bounds to its cognate BSs in CRE1, it forms a complex with bZIPs bound to CRM3. This protein interaction “zips” the tails of the loop which locks this conformation. Subsequently, the CRM1-CRM3 modules take over the transcriptional control of *Glu-1* genes from the basal promoter.

**Figure 6.**
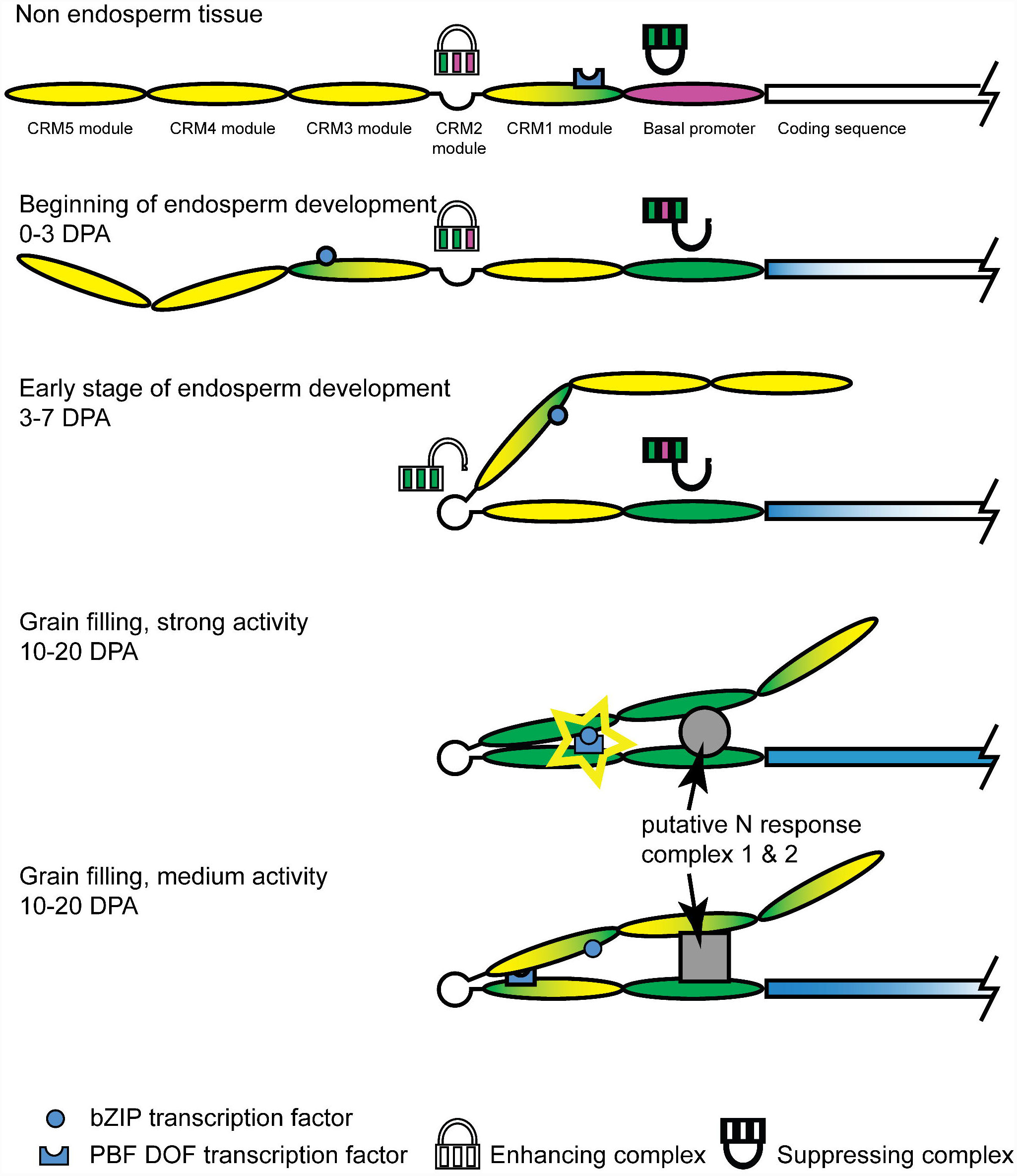
The model of the regulation of *Glu-1* genes. Modules of promoters are shown in a suppressed state in non-endosperm tissue and at different stage of endosperm development. The definitions of CRMs and basal promoter area are shown in Figure 4. Colour of modules shows their status: magenta – inhibition; yellow – neutral; green – driving expression; gradient – partially driving expression. Icons of combination lock represent transcription factor complexes. Each lock, has three wheels that represent TF families, the positions of the wheel represent various TFs of the same family. Green wheel indicates “correct position” of the wheel, aka appropriate TF is bound; magenta wheel indicates “wrong position”, aka wrong TF is bound or TF is missing. In case of suppressing complexes, the lock works in an inverse way, it opens when wrong combination is set, aka stops suppressing. The interaction of bZIP and PBF DOF TFs is shown for its enhancing effect. The colour intensity of blue in the coding sequences represents transcription activity. Grey circle and rectangle represents TF complexes of the two separate pathways of x and y-type *Glu-1* genes, respectively. These complexes are specified by BSs in the CRM4 and the basal promoter regions (see Figure 4.).

The temporarily fall back of activity for x-types genes in response to added nitrogen is at the same time than the event of loop zipping. Also at this time, there is a transcriptional reprogramming in developing seed as reported earlier (57–59). A change in the activity and/or in the composition of TFs caused by an nitrogen responsive element delays the loop formation and temporarily halt the activity compared to endosperm of the control plants.

Differential responses to environmental conditions of x-and y-type genes are coded in the CRM4, CRM5 and CRM6 and in the basal promoter regions. Differences in promoter profiles and expression profiles suggest two distinct regulatory mechanisms at action. In case of rice, it has been proposed that multiple regulatory mechanisms may be involved in the endosperm specific expression of glutelin genes (36). Earlier studies already reported that homoeolog wheat *Glu-1* genes are controlled by the same regulatory systems (60). Our findings are also in harmony with the general view that duplication is usually followed by divergence of expression and/or sub-functionalization (61).

On the other hand, different expression kinetics of genes is coded in the number of bZIP and DOF BSs in CRM3 and CRM1, respectively, as described above. However, differences in the activity of a transcription factor can also influence the activity of the target gene. It has been already proven that polymorphisms in the promoter regions of a bZIP TFs involved in grain filling has a direct result on wheat dough quality (62). Similarly, in barley, the differential activity of PBF DOF was reported to relate to grain protein content (63). In the case of rice, it has been demonstrated that DOF is not necessary for basic transcription and mostly acts as an enhancer and in certain construction as a suppressor (64). In cv. Glenlea, the early emergence of DOF transcription factors starts the full-blown transcription mechanisms as early as 5 DPA contrary to cv. Chinese Spring.

Possible further evidence of the interaction of distal regions of *Glu-1* promoters is the conserved, complementary pairs of MYB BSs in the basal promoter region and at the CRM4 region at equal distance from the CEREAL box. Although, this model does not specify the exact role of CEREAL-box in *Glu-1* promoters, it is assumed by its position that its presence influences the kinetics of loop formation. Based on the above regulatory model, the strong activity of *Glu-1Bx* genes can be interpreted by (i) an extra CEREAL-box facilitating DNA bending, (ii) an extra MYB BS present at CRM2 (position -548 on antisense) that enhances formation of the “looping” complex and (iii) the fact that there are no self-competing DOF BSs in CRM1.

Experiments with chimeric promoters with an *act*1 intron downstream of the basal promoter region of *Glu-1* genes increased the activity of reporter genes (65, 66). With regards of the proposed model, the intron of an ubiquitous gene acted as a direct enhancer to the basal promoter region. The above regulatory model does not require tissue specific transcription factors but rather a tissue specific group of transcription factors because inhibition, looping and “loop zipping” of bZIP/DOF interaction occurs via combinatorial effects.

## Conclusion

The function of promoter regions is to read out the transcription factor pool available in any given tissue at any given developmental stage. Different promoters are tuned to different factors. Never the less, similar promoters are tuned to similar factors and are assumed to produce similar expression dynamics if and only if the TF pools are identical. The facts that (i) promoter sequences of different *Glu-1* alleles are nearly identical; (ii) x-type and y-type *Glu-1* respond differently to environmental changes; (iii) promoter profiles are more related across homoeologs than paralogs and (iv) there are characteristic differences between *Glu-3* and *Glu-1* gene promoters; suggest that (a) the genes of LMW and HMW glutenin subunits do not share all aspects of their regulation, (b) there are at least two regulatory mechanism in action controlling *Glu-1* expression and (c) allelic differences in gene activity are caused by differences in transcription factor activity across genotypes. Therefore, our study concludes that the emergence of variation identified in the promoter profiles precedes the evolution of hexaploid wheat, which suggests that breeding had no influence on the polymorphism of *Glu-1* gene promoters and thereby on the amount of expressed HMW glutenin subunits. However, breeding could influence the expressed amount and the activity of the available transcription factors that are directly involved in the expression of *Glu-1* genes.

## SUPPLEMENTARY DATA

Supplementary Data are available at NAR online.

## ACKNOWLEDGMENTS

The authors are grateful to László Tora and Bob Anderssen for their careful reading, valuable discussions and their lot of useful advice. In addition, we would like to thank the International Wheat Genome Sequencing Consortium (IWGSC, www.wheatgenome.org) for pre-publication access to the IWGSC Chromosome Survey Sequences hosted at http://wheat-urgi.versailles.inra.fr/Seq-Repository.

